# Melatonin improves neuro-behavioral perturbations in diet/photoperiod induced chronodisruption

**DOI:** 10.64898/2026.03.09.710494

**Authors:** Aliasgar Vohra, Rhydham Karnik, Hitarthi Vyas, Shruti Kulshrestha, Whidul Hasan, Kapil Kumar Upadhyay, Helly Shah, Ranjitsinh Devkar

**Affiliations:** Division of Chronobiology and Metabolic Endocrinology, Department of Zoology, Faculty of Science, The Maharaja Sayajirao University of Baroda, Vadodara, Gujarat 390002, India; Dr. Vikram Sarabhai Institute of Cell and Molecular Biology, Department of Zoology, Faculty of Science, The Maharaja Sayajirao University of Baroda, Vadodara, Gujarat 390002, India; Department of Neurology, School of Medicine, Washington University in St Louis, St. Louis, MO, USA; Department of Internal Medicine, University of Michigan Medical School, Ann Arbor, MI, USA; Beth Israel Deaconess Medical Center (BIDMC), Harvard Medical, School, Harvard University, Boston, MA, USA

**Author notes:** Corresponding author full contact details. (1) Name: Ranjitsinh Devkar Address: Department of Zoology, The Maharaja Sayajirao University of Vadodara, Gujarat, India.

**Keywords:** Chronodisruption, High-fat-high-fructose-diet, Neurobehavior, Anxiety, Depression, Melatonin

## Abstract

Endogenous circadian oscillators regulate learning, cognitive performance and memory are disrupted due to circadian shifts. High-fat-high-fructose (H) diet, photoperiodic shifts induced chronodisruption (CD) and a combination (HCD) causes neurobehavioral perturbations wherein; the merits of exogenous melatonin in alleviating the said behavioral deficits are studied herein. Indices of anxiety (marble burying test, elevated plus maze test and hole board test) and depressive behavior (sucrose preference test, forced swim test and tail suspension test) were elevated in H, CD and HCD groups. Significant increments in the titres of thyroid hormone levels (T3, T4 and TSH) and mRNA levels of hippocampal pro-inflammatory genes (*Tnf-*α, *Il-1*β, *Il-4*, *Il-6*, *Il-10*, *Il-12*, *Il-17*, *Mcp-1* and *Nf-*κ*b*) in the said experimental groups corroborates with the said changes. Exogenous melatonin treatment to the said experimental groups viz. HM, CDM and HCDM; accounted for moderate to significant improvement in the said neurobehavioral perturbations and hippocampal inflammatory markers. Hippocampal BDNF-TrkB pathway genes of H, CD and HCD had recorded a non-significant downregulation in mRNA but without prominent changes in proteins. Likewise, melatonin-treated groups showed moderate to significant improvement in transcripts of *Bdnf, Trkb, Nt-3, Nt-4, Psd-95* and *Syn-1.* Herein, we report neurobehavioral perturbations caused by a combination of H and CD. Melatonin-mediated improvement in neurobehavior and the corrective changes in hippocampal BDNF-TrkB pathway implies towards the potential anxiolytic and anti-depressive activity as reported herein.

## Introduction

External factors such as diet, regular exercise, the feeding-fasting window and sleep-wake cycles contribute significantly to the overall well-being. Chronic consumption of diet rich in saturated fats and refined sugars is a major contributor of lifestyle disorders such as obesity and metabolic dysfunction-associated steatotic liver disease (MASLD)[1], as well as cognitive decline and dementia[2]. Hippocampus, located deep within the temporal lobe of the brain, is vital to learning and memory in mammals.[3] Mice subjected to high-calorie diets recorded a significant decrement in BDNF expression and impaired hippocampal-dependent memory, thereby resulting in decreased neuronal plasticity and deficits in learning and memory[4]. Further, depression and anxiety are often observed in MASLD patients.

Studies have reported that MASH patients, in particular, exhibit a significantly higher prevalence of major depressive disorder (MDD) and generalized anxiety disorder (GAD). [5], [6] Circadian desynchrony or chronodisruption (CD) due to exposure to artificial light at night, intentional sleep deprivation or consumption of high calorie diet is linked to metabolic disorders and behavioral deficits [7], [8], [9], [10], [11], [12]. Studies have shown that C57BL/6 mice subjected to aberrant light-dark (LD) conditions recorded increased corticosterone levels [13]. Adult mice subjected to a short photoperiod regime recorded alterations in brain size, hippocampal dendritic morphology and impaired long-term spatial learning and memory [14]. Growing evidence suggests that an altered photoperiod and high-calorie diets decreases hippocampal volume, impairs learning and cognitive function, diminishes recognition memory and causes locomotor deficits as well as impairs learning and attention in humans. Notably, these alterations in photoperiod and diet also increase the risk of depression and anxiety.

Melatonin, a neuro-hormone (N-acetyl-5methoxytryptamine) is known to improve hepatic lipid accumulation in high-fat diet-fed (HFD) mice [15]. Antioxidant and cytoprotective properties of melatonin are known to reverse the low degree of inflammatory tissue damages in experimental models of neurodegenerative disorders, aging, Alzheimer’s, and Parkinson’s disease [16], [17], [18]. Melatonin is a well-established neuroprotective regulator that counters neurotoxicity by stabilizing mitochondrial function and activating AMPK-driven mitophagy, thereby strengthening resilience against circadian and metabolic stress.[19] Further, an improvement in oxidative stress, neuro-inflammation and BDNF-TRKB pathway in hippocampus of rats following melatonin treatment has been accredited to an improved behavioral pattern [20]. Fluoxetine-treated (higher anxiety and depression) male C57BL/6N mice were reported for an improvement in BDNF-TrkB pathway in hippocampus following melatonin treatment [21], [22]. Further, exogenous melatonin can improve cognition through neuroprotective mechanism against HFD-induced neurobehavioral stress but, the underlying mechanism for the same lacks clarity.

The underlying mechanisms of neurobehavioral perturbations due to a combination of chronodisruption (CD) and high-fat-high-fructose (H) diet lacks clarity. Further, studies that evaluate the corrective effect of melatonin in the said experimental regimen are lacking. Herein, we assess the merits of melatonin in alleviating hippocampal pro-inflammatory changes in CD and/or H treated C57BL/6J mice. Additionally, we report on melatonin-mediated improvements in thyroid hormone levels, hippocampal BDNF-TrkB pathway intermediates and the consequent corrective changes in behavioral disturbances.

## Experimental procedures

### Chemicals and Reagents

TRIzol, and SYBR green (SYBR select master mix) was procured from Invitrogen (Thermo Fisher Scientific, USA). Reagents for protein assay, Polyvinylidene fluoride (PVDF) membrane, Precision Plus protein ladder, iScript cDNA synthesis kit and Clarity Western ECL substrate were procured from Bio-Rad Laboratories (CA, USA). Antibodies against viz. BDNF (E-AB-18244), ERK1/2 (E-AB-12397), and SYN-1 (E-AB-33003) (Everon Life Sciences), β-actin (4970S) and anti-rabbit secondary antibody (7074P2) (Cell Signalling Technology, MA, USA) and TrkB (MAB397-SP) (Biotechne, MN, USA). Other chemicals such as RNA Later (Ambion Inc., Thermo Fisher Scientific, USA). Thyroid ELISA kit (T_3_, T_4_and TSH; XEMA-MEDICA Co., Ltd. Russia), Melatonin and Protease inhibitor cocktail (PIC; Sigma Aldrich MO, USA) and the miscellaneous purchases from Merck (Darmstadt, Germany).

### Animal studies

C57BL/6J(Male) mice (60, 6-7 wk, 20-22g) were procured from ACTREC, Mumbai, India and maintained as per CCSEA standard guidelines (23±2°C, LD 12:12, laboratory chow and water ad libitum). All experimental protocols were reviewed and approved by the Institutional Animal Ethics Committee (IAEC; Approval No. MSU/Z/IAEC03/01-2019) and conducted in a CPCSEA-approved animal facility (827/GO/Re/S/04/CPCSEA), Department of Zoology, The Maharaja Sayajirao University of Baroda. Experiments were performed in compliance with ARRIVE guidelines. Mice were divided randomly (n=8 per group x 7 groups) and acclimatized for a week. Control mice were fed with laboratory chow whereas the experimental mice were fed high-fat-high-fructose (H) diet and/or subjected to chronodisruption. Daily evening melatonin injections were started from 10^th^ to 18^th^ week.[23], [24]

### Photoperiodic/dietary regimens and experimental manipulations

Experimental groups: (i) control, (ii) CD (chow diet and CD –chronodisruption), (iii) H (high-fat-high-fructose diet + 20% fructose water) and (iv) H + CD and maintained for 18 weeks, (v) CDM (CD + Melatonin), (vi) HM (H + Melatonin and (vii) HCDM (H + CD + Melatonin). Groups (v), (vi) and (vii) were daily injected (*ip*) with melatonin (10mg/kg) after 8 weeks as per Joshi et al. 2021[24]. Chronodisruption (CD) was induced by phase-advance-phase-delay regimen, that is a standardized protocol of our laboratory [4], [23], [24], [25], [26]. Briefly, mice were housed in two different rooms: Room 1 (7:00 h to 19:00 h; light period/photophase and 19:00 h to 7:00 h; dark period/scotophase) and Room 2 (11:00 h to 23:00 h; dark period/scotophase and 23:00 h to 11:00 h; light period/photophase). Mice were shifted from Room 1 to Room 2 on Monday (lights off at ZT4; phase advance of 8 h) and transferred back from Room 2 to Room 1 on Thursday (lights off at ZT20; phase delay of 8 h). All the transfers were carried out at 10:55 h. Animal health and welfare were monitored throughout the study by trained personnel. Cage-side observations were conducted on alternate days, and more frequently when required. Parameters monitored included: body weight changes (>20% loss), food and water intake, grooming behavior, posture and locomotor activity, fur condition, signs of dehydration or lethargy, abnormal aggression or stereotypic behavior[4], [24].

### Humane endpoints and termination of experiment

Animals were predefined to be humanely euthanized if any of the following criteria were observed: body weight loss exceeding 20% of baseline, persistent hypoactivity or inability to access food/water, severe lethargy, hunched posture, or unresponsiveness, signs of pain or distress unrelieved by supportive care. Once humane endpoint criteria were met, animals were scheduled for euthanasia. Importantly, death was not a planned experimental endpoint, and no animals died prior to reaching humane endpoint criteria during the course of the study. At the end of the 18-week experimental period, animals were anesthetized using isoflurane. Blood was collected via retro-orbital sinus puncture under anesthesia, followed by euthanasia by anesthetic overdose in accordance with IAEC and CPCSEA guidelines. Brain tissues were harvested immediately and stored either in RNAlater for gene expression analysis or at −80 °C for immunoblot studies. All procedures were performed with maximal efforts to minimize animal discomfort, pain, and distress.[15], [23], [27]

### Serum parameters

Liver function markers (AST, ALT and ALP; Reckon Diagnostic Ltd., India), glucose and creatinine were estimated.

### Serum thyroid hormones analysis by ELISA

ELISA of Thyroid hormones (T_3_, T_4_ and TSH) in serum titres (XEMA-MEDICA Co., Ltd. Russia) was done following manufacturer’s protocol. [28]

### Behavioral tests

The behavioral tests for control and experimental groups of mice were performed during ZT3 to ZT9 (10-00 a.m. to 4-00 p.m.). Sterile conditions were maintained for apparatus of training/probe trials and olfactory or spatial clues were eliminated from the testing area for all groups. Background white noise (of < 60 db and zero vibrations) was used during the tests. Mice were acclimatized for an hour and subsequently, behavioral tests viz., hole board, marble burying and elevated plus maze test (for anxiety) and force swim and tail suspension test (for depression-like behavior) were performed. High resolution videos of behavioral test were recorded and data was analysed using ANYMaze software (Stoelting Co., Wood Dale, IL).

### Force swim test (FST; for depression)

Immobility time in mice was evaluated in an open cylindrical container (dia. 10 cm, ht. 25 cm), containing 19 cm water (25 ± 1 °C). Each mouse was video graphed and the calculations of mobile time (desperate swimming and an effort to come out of the container) or immobile time (when mice ceased to struggle and remained motionless on water surface) was noted. High indices of immobility time indicate a depression-like activity.[29]

### Tail suspension test (TST; for depression)

In TST, adhesive tape was used to suspend the mice to a horizontal bar for 6 minutes and video-graphed. Classically, suspended mice immediately perform escape-like behavior or an immobile posture. Prolonged periods of immobility were considered to be an index of depression-like behavior. Video clips were visually analysed to record immobility time (by a volunteer blinded to the experimental groups).

### Marble-burying test (MBT; for anxiety)

Mice were segregated in experimental cages (4–5 cm-thick rice husk) for 10 min (habituation phase) followed by a 40-minutes home cage resting period. Later, mice were re-introduced into the experimental cages (for 30 min), comprising of twenty sterilized glass marbles (diameter[10 mm; overlaid on the bedding) arranged in a 4×5 matrix. The cages were photographed at 0, 10, 20 and at 30 min and the marbles were re-counted.

### Hole-board test (HBT; for anxiety)

The hole-board test (HBT) relies on the modulation of natural exploratory behavior, wherein stress inhibits exploration, and anxiolytic treatments counteract this effect. Herein, the numbers of head dips in holes on a board in an infrared actimeter as mentioned above. The floor is made of opaque Plexiglas with evenly distributed 16 holes (each 3.8 cm diameter and 10 cm deep). Total counts for the head-dips were recorded from the digital panel of the actimeter.

### Elevated plus maze test (EPM, for anxiety)

The elevated plus maze was used during the experiment had two closed arms (5×30 cm, with a clear 15-cm-high perspex walls) and two open arms (5×30 cm long) placed at a height of 40 cm. In a single 5-min trial, number of entries of mice in each arm, and the time spent (in sec) was analysed recorded by a camera after being placed at the centre of the maze and the data was analysed using ANYMaze Video Tracking System (Stoelting, Wood Dale, IL, USA).

### Gene expression studies by qPCR

TRIzol based extraction of total RNA from hippocampal tissue was done and cDNA was made (iScript cDNA Synthesis kit). mRNA of various genes was quantified by qPCR analysis (QuantStudio 3, Life Technologies, CA, USA) using SYBR green SYBR Select Master Mix. All the data was normalized with GAPDH and analysed using the 2^−ΔΔCT^ method. Gene-specific mice primers used for this study are listed in Supplementary Table 1.[26]

### Immunoblots analysis

Hippocampal tissue was homogenized in RIPA buffer containing PIC and 1 mM PMSF followed by incubation in ice-bath for 2h as previously described in Karnik et al. 2024 [4]. The tissue lysates were spinned (12,000 rpm at 4°C for 20 min) and protein was estimated in supernatant (Bio-Rad protein assay reagent kit, USA). Protein samples were denatured (6X loading dye at 95°C-100°C for 5 min) and 25 μg was separated by SDS-PAGE and subsequently transferred onto PVDF membrane (Trans-Blot Turbo Transfer System, Bio-Rad Laboratories, USA). Transfer of proteins was checked by staining membrane with 0.05% Ponceau S. Membrane was de-stained in distilled water followed by blocking with 3% BSA in Tris buffered saline (TBS) for 1 h at RT. Subsequently, membrane was incubated overnight in anti-BDNF (1:1000), anti-TrkB (1:1000), anti-ERK1/2 (1:1000) and anti-SYN-1 (1:1000) antibodies in 3% BSA. The membrane was subjected to three TBS washes with 0.1% Tween 20 (TBST) and probed with HRP-linked anti-rabbit secondary antibody (1:5000) for 1 h at RT. Blots were developed (Clarity western ECL reagent, Bio-Rad, CA, USA) and resolved using X-ray films. Anti-β-actin antibody (1:5000) was used to determine equivalent loading.

### Statistical analysis

Data (mean ± SD) was further analysed statistically with one-way analysis of variance (ANOVA) and Tukey’s multiple comparison tests using Graph Pad Prism 5.0 (CA, USA) in comparison with control and disease control. *p< 0.05, **p< 0.01 and ***p< 0.001 were significant as compared to control. #p< 0.05, ##p< 0.01 and ###p< 0.001 were significant when melatonin-treated groups (CDM, HM and HCDM) were compared with the respective disease-control (CD, H and HCD) groups.

## Results

### Melatonin mediated changes in body, liver and adipose tissue weights

At the end of the experimental period, the body, liver and adipose weights and body circumference of H (p<0.001) and HCD (p<0.01) groups were significantly higher than the control group. However, CD group did not record any significant change in the said parameters. The food and water intake were non-significantly lower in all the experimental groups throughout the period of study. Melatonin treatment accounted for a significant decrement in body weight and body circumference of HM (p<0.001) and HCDM (p<0.05) groups. The said parameters for CDM group were comparable to that of control. ( Supplementary Figure 1)

### Melatonin improves serum liver function markers, glucose and creatinine

Liver function markers (ALT, AST and ALP) were assayed in serum of control and treated mice. A significant increment (p< 0.001) was observed in the circulation titres of ALT and AST in CD, H, and HCD groups with non-significant changes in the ALP levels. Further, CD group recorded a non-significant decrement in serum glucose levels whereas H and HCD groups recorded a non-significant increment. Additionally, H, CD and HCD groups recorded a non-significant increment in the levels of serum creatinine. Melatonin treatment significantly lowered serum ALT and AST levels in CDM, HM, and HCDM groups. Melatonin-treated groups also recorded a non-significant increment in glucose levels whereas; a significant decrement was observed in HM (p<0.01) and HCDM (p<0.001) groups. Furthermore, melatonin accounted for a non-significant improvement in serum creatine in HM and HCDM groups and a significant improvement in CDM (p<0.01) group. (Figure 1)

**Figure 1.**
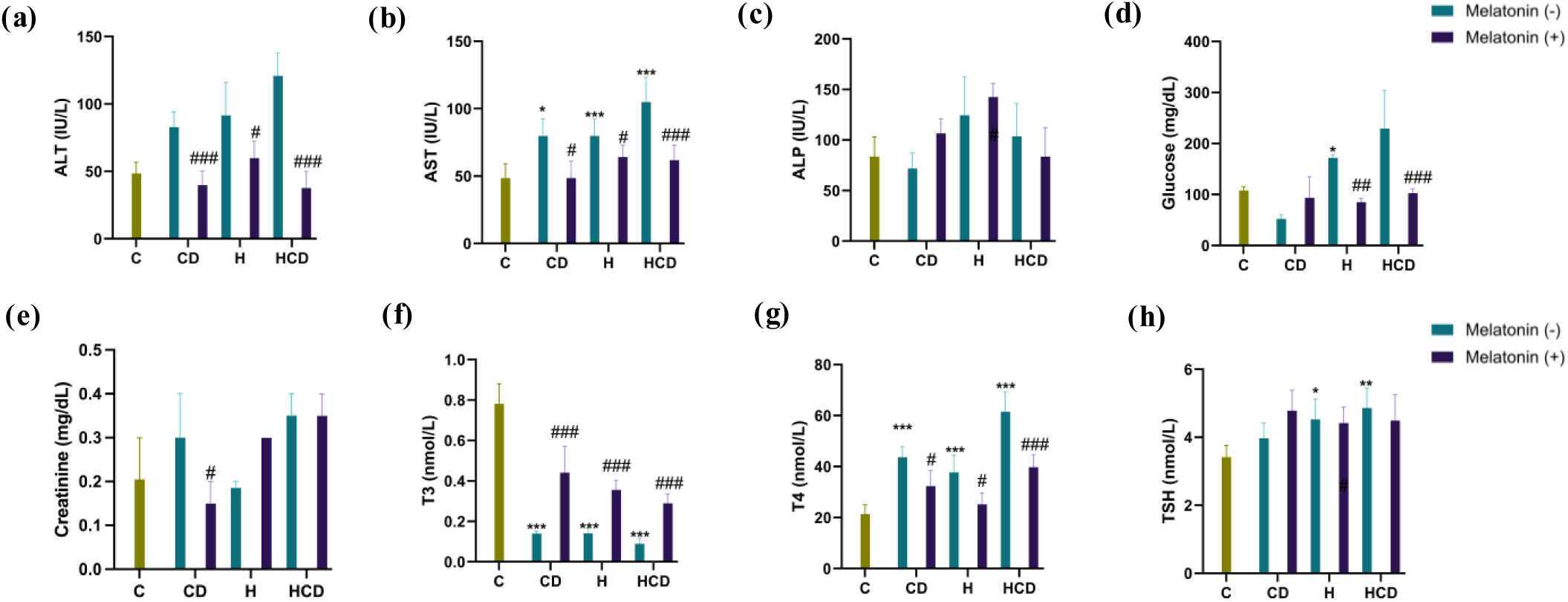
Serum liver function markers. (a) ALT (IU/L) (b) AST (IU/L) (c) ALP (IU/L) (d) Glucose (IU/L) and (e) Creatinine (IU/L). Thyroid hormone titers (f) T3 (nmol/L) (g) T4 (nmol/L) and (h) TSH (nmol/L). Significant at p-value (< 0.05); (one-way ANOVA test followed by Bonferroni’s Multiple Comparison Test), values expressed as mean ± SD, (n = 6) *p < 0.05, and ***p < 0.001 is when CD, H and HCD compared to Control (C). #p < 0.05, ##p < 0.01, and ###p < 0.001 is when CDM compared with CD, HM with H and HCDM with HCD respectively.

### Melatonin treatment improves thyroid hormones levels in H and/or CD mice

Circulating titres of serum T_3_ were significantly lowered (p<0.001) whereas; T_4_ and Thyroid Stimulating Hormone (TSH) were found to be elevated (p<0.001) in CD, H and HCD groups. Melatonin treated groups recorded a significant increment (p<0.001) in serum T_3_ levels and a decrement in serum T_4_ levels in CDM, HM and HCDM groups as compared to their respective disease control groups but did not record any significant changes in TSH levels in melatonin-treated groups. (Figure 1)

### Melatonin lowers hippocampal inflammation in CD, H and HCD groups

In hippocampal tissue, mRNA levels of pro-inflammatory (TNF-α, MCP-1, NFκB, IL-1β, IL-6, IL-12, IL-17, CREB and IBA-1) and anti-inflammatory (IL-4 and IL-10) cytokines markers were assessed. Moderate to significant increments were recorded in the mRNA levels of pro-inflammatory cytokines, in H (NFκB, IL-1β & IL-6), CD (IL-1β & CREB) and in HCD groups (IL-12 & CREB). Melatonin-mediated lowering of inflammation was evident in form of significant decrement in mRNA levels in HM (NFκB, IL-1β & IL-6), CDM (IL-6 & CREB) and HCDM (NFκB) groups. Other pro-inflammatory marker genes (TNF-α, MCP-1, IL-12, IL-17, CREB and IBA-1) recorded a non-significant decrement following melatonin treatment. Melatonin treatment also accounted for significant improvement in mRNA levels of anti-inflammatory cytokines IL-4 levels in CDM and HCDM groups, whereas IL-10 showed a non-significant increment following melatonin treatment. (Figure 2)

**Figure 2:**
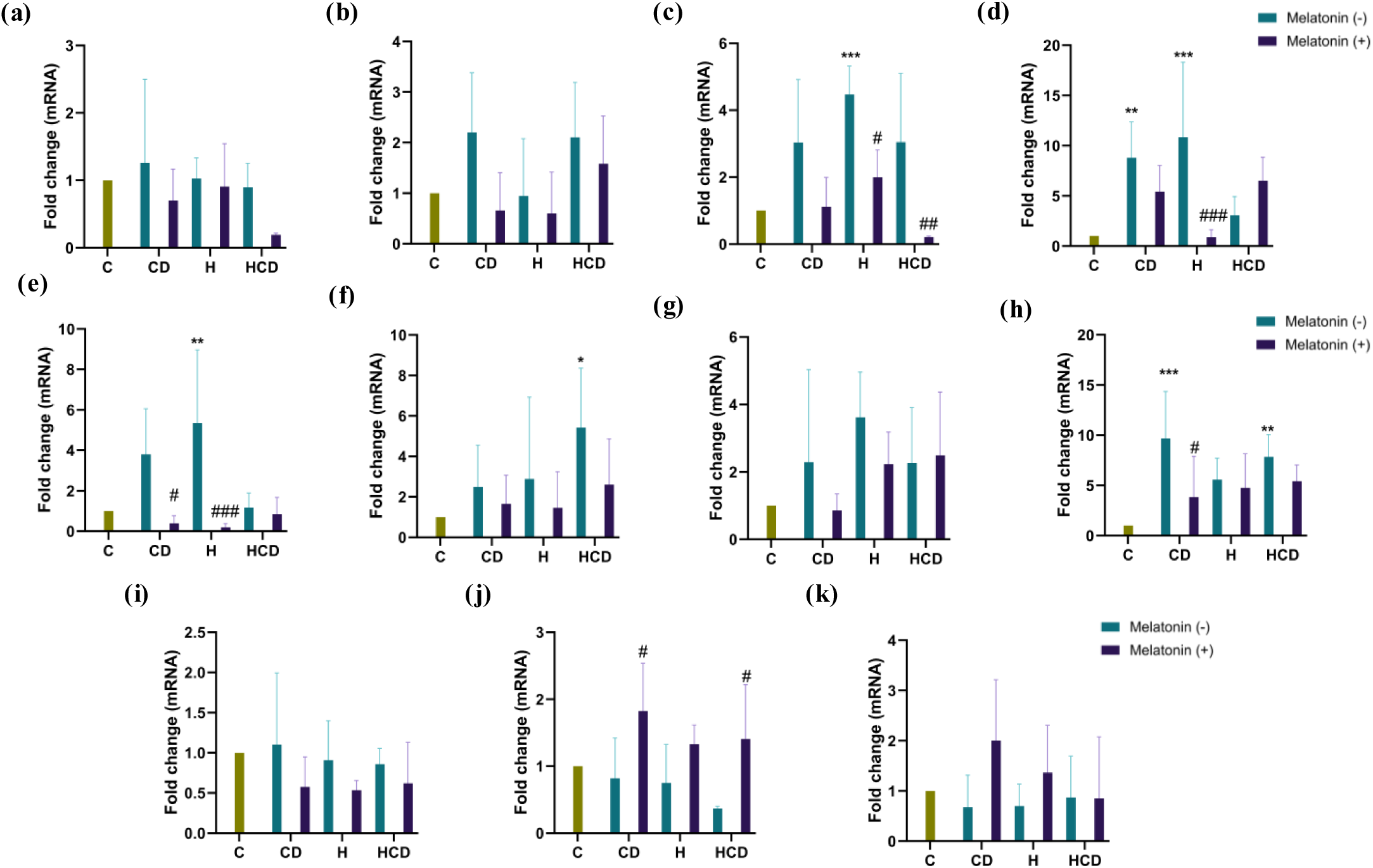
Hippocampal inflammatory markers. The mRNA levels of (a) TNF-α (b) MCP-1(c) NF-κB (d) IL-1β (e) IL-6 (f) IL-12 (g) IL-17 (h) CREB (i) IBA-1 (j) IL-4 (k) IL-10. Significant at p-value (< 0.05); (one-way ANOVA test followed by Bonferroni’s Multiple Comparison Test), values expressed as mean ± SD, (n = 6) *p < 0.05, and ***p < 0.001 is when CD, H and HCD compared to Control (C). #p < 0.05, ##p < 0.01, and ###p < 0.001 is when CDM compared with CD, HM with H and HCDM with HCD respectively.

### Melatonin mediated re-wiring of behavioural deficits Tests for depression

Sucrose preference test (SPT) is a reward-based test that is used as an indicator of decreased ability to experience pleasure or anhedonia and the same was found to be significantly lower in CD mice (p<0.01). Other experimental groups recorded a non-significant increment in the said parameter. ( Figure 3a)

**Figure 3.**
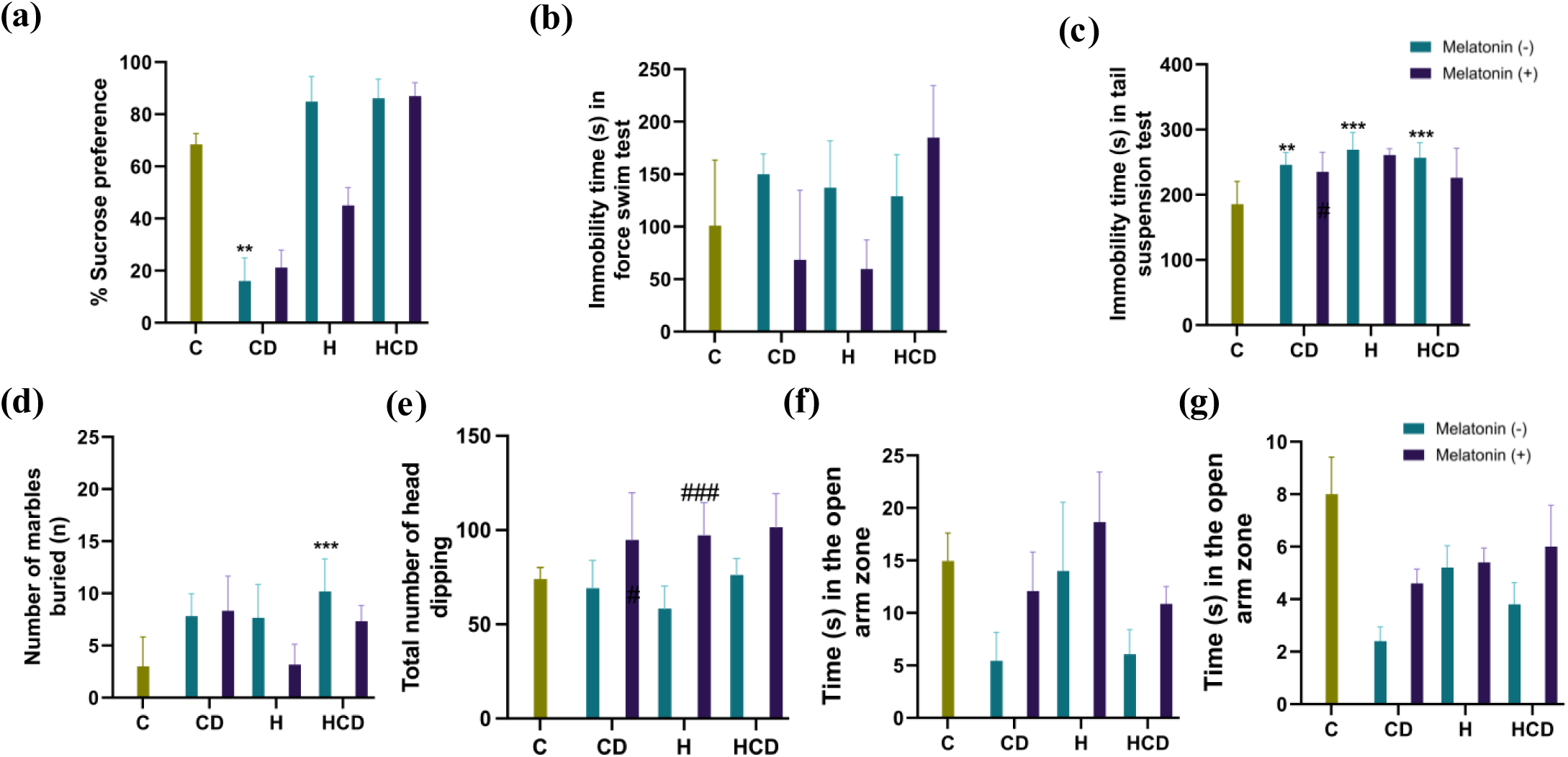
Melatonin mediated re-wiring of behavioural deficits. Assessment of Depression-type phenotypes using (a) Sucrose preference test (b) Force Swim Test (FST) and (c) Tail Suspension Test (TST). Assessment of Anxiety-type phenotypes using (d) Marble Burying Test (MBT) (e) Hole Board Test (HBT) and Elevated Plus Maze (EPM) test (f) Time spent in open arm zone (g) Number of enteries in open arm zone. Significant at p-value (< 0.05); (one-way ANOVA test followed by Bonferroni’s Multiple Comparison Test), values expressed as mean ± SD, (n = 6) *p < 0.05, and ***p < 0.001 is when CD, H and HCD compared to Control (C). #p < 0.05, ##p < 0.01, and ###p < 0.001 is when CDM compared with CD, HM with H and HCDM with HCD respectively.

Force swim test (FST) and tail suspension test (TST) are commonly used to assessed depression like behaviour in rodent models. In FST and TST, a longer duration of immobility of mice accounts for depression like behaviour. In our study, the CD, H and HCD groups recorded non-significant increment in FST and a significant increment (p<0.01 and p<0.001) in TST as compared to control. Melatonin treatment accounted for a non-significant decrement in the immobility time in FST and TST.(Figure 3b and 3c)

### Tests for Anxiety

Anxiety-like behaviour in mice was assessed by marble burying (MBT), hole board (HBT) and elevated plus maze (EPM) tests. In MBT, higher number of buried marbles were considered as an indirect evidence of elevated anxiety wherein, CD, H and HCD (p<0.001) groups recorded moderate to significant increment in MBT indices with the one in HCD group being significantly higher than control (p<0.001). Melatonin treatment accounted for moderate decrement in HM and HCDM groups in both diurnal and nocturnal phases. (Figure 3d)

The hole board test (HBT) records the number of head dips in the hole wherein; CD, H and HCD recorded a moderate decrement in the number of head dipping. Melatonin treatment significantly increased the head dipping score in HM (p<0.01) group as compared to H group. Non-significant increments were observed in CDM and HCDM groups as compared to their respective disease control (CD & HCD) groups. (Figure 3e)

In the elevated plus maze test (EPM) the number of entries and time spent in the open arm of a plus maze is an index of lesser anxiety. In this study, CD, H and HCD groups showed moderately lower number of diurnal entries in open arm, implying towards anxiety-like behaviour. (Figure 3g) The time spent in the open arm of an EPM too showed a decrement in CD, H and HCD groups. Melatonin-treated groups revealed higher number of EPM entries in CDM and HCDM groups whereas; HM group recorded no significant change in the said parameter. (Figure 3f)

### Melatonin treatment favourably impacts the hippocampal BDNF-TrkB pathway genes

Hippocampal region in the brain of control and treated mice were assessed for possible changes in the key genes (BDNF, TrkB, Syn-1, & Psd95) of the *BDNF-TrkB* pathway. The mRNA levels of the said genes were found to be lowered in CD & H groups with significant improvement in melatonin treated. Similarly, decrements were recorded in mRNA levels of neurotrophin genes (Nt-3 & Nt-4) with melatonin treatment accounting for an improvement. (Figure 4a-f)

**Figure 4:**
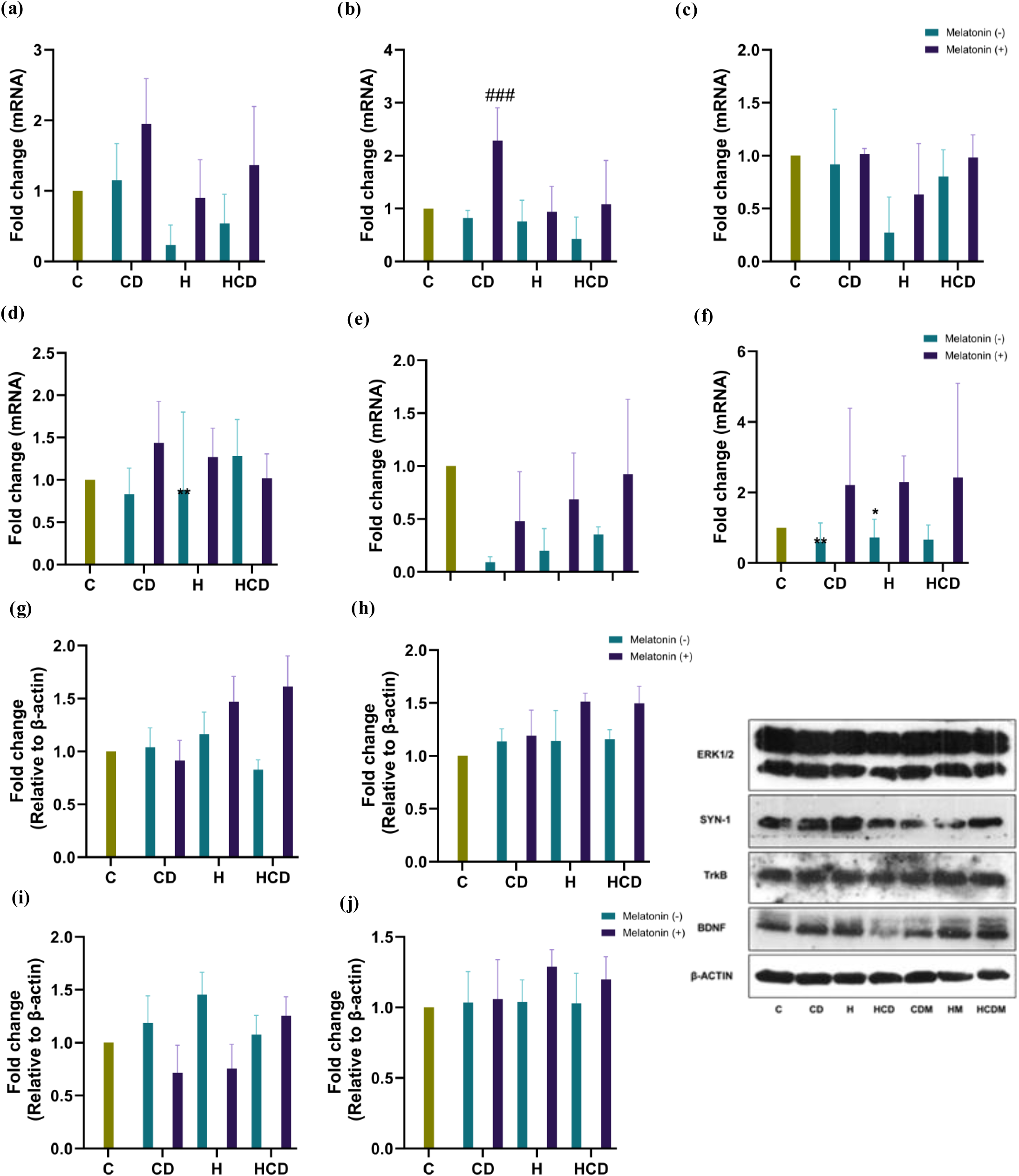
Melatonin impacts hippocampal BDNF-TRkB pathway. The mRNA levels of (a) BDNF (b) TRkB (c) SYN-1 (d) PSD-95 (e) NT-3 and (f) NT-4. Their respective protein levels (g) BDNF (h) TRkB (i) SYN-1 (j) ERK 1/2. Significant at p-value (< 0.05); (one-way ANOVA test followed by Bonferroni’s Multiple Comparison Test), values expressed as mean ± SD, (n = 6) *p < 0.05, and ***p < 0.001 is when CD, H and HCD compared to Control (C). #p < 0.05, ##p < 0.01, and ###p < 0.001 is when CDM compared with CD, HM with H and HCDM with HCD respectively.

Protein levels of BDNF recorded no change in CD and H groups whereas; HCD group recoded a moderate non-significant decrement. Melatonin treatment accounted for a moderate improvement in BDNF protein in HM and HCDM groups. (Figure 4g) Immunoblots of TrkB, SYN-1 and ERK1/2 in CD, H and HCD groups were comparable to that of control. Further, melatonin treatment did not account for any significant change in the protein expression of the said genes. (Figure 4h-j) Overall, the mRNA levels of BDNF-TrkB and neurotrophins genes showed a decrement in disease control groups (CD, H and HCD), but the same was not reflected in its protein levels.

## Discussion

Lifestyle disorders, particularly obesity, diabetes and NASH, are increasingly linked with mood alterations, anxiety, depression, and cognitive decline. These behavioural perturbations are often underpinned by circadian misalignment caused by aberrant photoperiods or synergistic dietary factors[4]. In the present study, we employed a model wherein C57BL/6J mice were subjected to photoperiod-induced chronodisruption (CD) and/or high-fat-high-fructose (H) diet alone; or in a combination (HCD). Herein, we demonstrate a potent therapeutic role of melatonin in rescuing neurobehavioral deficits by targeting hippocampal inflammation, oxidative stress, and neurotrophic signalling.

Initially, we confirmed hepatic and metabolic dysfunction (in CD, H, and HCD mice), consistent with our previous work (Joshi et al., 2021; Vohra et al., 2024). Crucially, we identified a concomitant disruption of the hypothalamic-pituitary-thyroid (HPT) axis, characterized by a reduction in circulating T3 and TSH alongside elevated T4[28], [30], [31]. The restoration of T3, T4, and TSH titres upon melatonin treatment aligns with growing evidence of melatonin’s direct beneficial effects on thyroid homeostasis in models of NASH and circadian desynchrony (Gładysz et al., 2023; Singh et al., 2022).

Given the established vulnerability of hippocampal neurons to metabolic insult (Miao et al., 2023), our findings revealed elevated pro-inflammatory cytokine transcripts in the hippocampus of CD, H, and HCD groups. Melatonin treatment effectively suppressed this neuroinflammatory response, an effect that can be attributed to the downregulation of NF-κB. This aligns also with the established paradigm wherein melatonin exerts potent anti-inflammatory effects, in part through the activation of SIRT1, which deacetylates and inhibits NF-κB, thereby reducing the transcription of inflammatory mediators (Keskin-Aktan et al., 2022; Luchetti et al., 2022). Interestingly, the magnitude of melatonin’s corrective effect varied depending on the insult, with HCDM animals showing only partial recovery in depressive-like behaviour. This suggests that while melatonin is efficacious against isolated stressors, its ability to counteract compounded dietary and circadian insults may require optimization in dose regimen and timing of administration.

The consequences of neuroinflammatory milieu was quantified through a battery of behavioral tests. In the forced swim test (FST), a measure of depressive-like behavior, melatonin significantly reduced immobility time in the CDM and HM groups. This antidepressant-like effect is consistent with reports that melatonin activates the Nrf2 pathway, counteracting oxidative stress and inflammation induced by insults like LPS (Qin et al., 2022). Behavioural outcomes in anxiety paradigms (elevated plus maze, marble burying and hole board) demonstrated that melatonin consistently improved exploratory behaviour and reduced anxiety-like indices. These findings reinforce melatonin’s anxiolytic role, and align with reports of improved locomotor performance in neuroinflammatory models [4]. The antidepressant effect, however, appeared less robust under combined diet and photoperiod stress, pointing towards mechanistic differences in how melatonin modulates anxiety versus depression-like circuits.

The H group had recorded a significant downregulation of BDNF, Synapsin-1 (Syn-1), and NT-3 suggesting a diet-dominant effect on the BDNF-TrkB pathway and a direct link between metabolic dysregulation and impaired synaptic resilience. Key role of melatonin as a regulator of neuroplasticity, capable of counteracting the synaptotoxic environment created by poor diet and erratic light cycles. This pro-neurotrophic action provides a compelling mechanistic foundation for the observed improvements in behavioral flexibility and cognitive health. These findings are consistent with earlier reports of melatonin augmenting hippocampal neurotrophins under high-fat diet or sleep-deprivation paradigms.

Taken together, our study provides compelling evidence that combined chronodisruption and high-fat-high-fructose diet synergistically disrupt behavioural and molecular homeostasis. Melatonin mitigates these effects by modulating thyroid hormone balance, suppressing hippocampal inflammation, and enhancing neurotrophic signalling. While anxiolytic benefits were consistent, antidepressant outcomes appeared contingent upon the severity of combined insults. Our findings delineate a multifaceted pathway whereby diet and photoperiod-induced chronodisruption converge to drive neurobehavioral pathology through systemic endocrine disruption, hippocampal neuroinflammation, and a deficit in neurotrophic signalling. The present findings not only add to the mechanistic understanding of melatonin’s neuroprotective role, but also highlight its potential clinical relevance in patients suffering prolonged with lifestyle disorders associated with circadian disruption.

## Conclusion

A combination of dietary and photoperiodic shifts induced chronodisruption (HCD group) provides a peep in the sojourn of lifestyle disorder that culminates in behavioural perturbations. Exogenous melatonin administration provides evidence of improving behavioural deficits in condition of lifestyle disorder (such as MASLD) by improving: (a) circulating titres of thyroid hormones, (b) hippocampal inflammation and (c) anxiety and depression-like behaviour. Overall, the outcomes provide evidence on role of melatonin in modulating neurobehavior; a finding that can improve the mental health of patients suffering from prolonged durations of lifestyle disorders.

## Acknowledgements

This work was supported by the Department of Biotechnology, Government of India (DBT: BT/PR36169/MED/30/2197/2020).

## Author contributions

A.V., and R.D. designed the study; R.D. obtained funding; in vivo studies and molecular biology experiments were performed by A.V., K.U., H.V., R.K. and S.K.; behavioral data was analysed by A.V., K.U., and W.H.; the manuscript was written by A.V., S.K. and H.S.; R. D. analysed the data, contributed, and edited manuscript. All the authors reviewed the final manuscript.

## References

[1] P. Jegatheesan and J. P. De Bandt, “Fructose and NAFLD: The multifaceted aspects of fructose metabolism,” Nutrients, vol. 9, no. 3, Mar. 2017, doi: 10.3390/NU9030230.

[2] R. Agrawal and F. Gomez-Pinilla, “‘Metabolic syndrome’ in the brain: deficiency in omega-3 fatty acid exacerbates dysfunctions in insulin receptor signalling and cognition,” J Physiol, vol. 590, no. 10, pp. 2485–2499, May 2012, doi: 10.1113/JPHYSIOL.2012.230078.

[3] Y. Miao et al., “The Presence and Severity of NAFLD are Associated With Cognitive Impairment and Hippocampal Damage,” J Clin Endocrinol Metab, vol. 108, no. 12, pp. 3239–3249, Nov. 2023, doi: 10.1210/CLINEM/DGAD352.

[4] R. Karnik et al., “Diet/photoperiod mediated changes in cerebellar clock genes causes locomotor shifts and imperative changes in BDNF-TrkB pathway,” Neurosci Lett, vol. 835, p. 137843, Jul. 2024, doi: 10.1016/J.NEULET.2024.137843.

[5] I. M. Balmus et al., “Neurological and neuropsychiatric comorbidities occurring in fatty liver diseases,” European Psychiatry, vol. 66, no. Suppl 1, p. S920, 2023, doi: 10.1192/J.EURPSY.2023.1945.

[6] C. D. Moulton, J. C. Pickup, and K. Ismail, “The link between depression and diabetes: The search for shared mechanisms,” Lancet Diabetes Endocrinol, vol. 3, no. 6, pp. 461–471, Jun. 2015, doi: 10.1016/S2213-8587(15)00134-5.

[7] R. Karnik, A. Vohra, H. Vyas, and R. Devkar, “Abstract 2026: Epigenetic and transcriptional regulation of CXCR4 by circadian clock regulated miRNAs in human monocyte derived macrophages,” Journal of Biological Chemistry, vol. 299, no. 3, p. 104382, Jan. 2023, doi: 10.1016/j.jbc.2023.104382.

[8] A. Joshi, K. K. Upadhyay, A. Vohra, K. Shirsath, and R. Devkar, “Melatonin induces Nrf2-HO-1 reprogramming and corrections in hepatic core clock oscillations in Non-alcoholic fatty liver disease,” The FASEB Journal, vol. 35, no. 9, pp. e21803–e21803, 2021.

[9] R. Karnik, A. Vohra, H. Vyas, N. Dalvi, S. Kulshrestha, and R. Devkar, “EP1219: MELATONIN IMPROVES GUT DYSBIOSIS IN HIGH FAT-HIGH FRUCTOSE DIET MODEL OF NAFLD AND/OR PHOTOPERIOD INDUCED CHRONODISRUPTION IN C57BL/6J MICE,” Gastroenterology, vol. 162, no. 7, p. S-1290–S-1291, May 2022, doi: 10.1016/s0016-5085(22)63780-6.

[10] A. Vohra et al., “Melatonin-mediated corrective changes in gut microbiota of experimentally chronodisrupted C57BL/6J mice,” Chronobiol Int, 2024, doi: 10.1080/07420528.2024.2329205.

[11] “Karnik: EP1219: MELATONIN IMPROVES GUT DYSBIOSIS… - Google Scholar.” Accessed: May 27, 2024. [Online]. Available: https://scholar.google.com/scholar?hl=en&as_sdt=0,5&cluster=305387985778487102 5

[12] J. Bass and M. A. Lazar, “Circadian time signatures of fitness and disease,” Science (1979), vol. 354, no. 6315, pp. 994–999, Nov. 2016, doi: 10.1126/SCIENCE.AAH4965.

[13] L. K. Fonken et al., “Light at night increases body mass by shifting the time of food intake,” Proc Natl Acad Sci U S A, vol. 107, no. 43, pp. 18664–18669, Oct. 2010, doi: 10.1073/PNAS.1008734107.

[14] L. M. Pyter, B. F. Reader, and R. J. Nelson, “Short photoperiods impair spatial learning and alter hippocampal dendritic morphology in adult male white-footed mice (Peromyscus leucopus),” Journal of Neuroscience, vol. 25, no. 18, pp. 4521–4526, May 2005, doi: 10.1523/JNEUROSCI.0795-05.2005.

[15] A. H. Vohra, K. K. Upadhyay, A. S. Joshi, H. S. Vyas, J. Thadani, and R. V. Devkar, “Melatonin-primed ADMSCs elicit an efficacious therapeutic response in improving high-fat diet induced non-alcoholic fatty liver disease in C57BL/6J mice,” Egyptian Liver Journal, vol. 11, no. 1, Dec. 2021, doi: 10.1186/S43066-021-00157-W.

[16] S. Pérez-Lloret and D. P. Cardinali, “Melatonin as a Chronobiotic and Cytoprotective Agent in Parkinson’s Disease,” Front Pharmacol, vol. 12, p. 650597, Apr. 2021, doi: 10.3389/FPHAR.2021.650597/XML.

[17] R. J. Reiter, J. C. Mayo, D. X. Tan, R. M. Sainz, M. Alatorre-Jimenez, and L. Qin, “Melatonin as an antioxidant: under promises but over delivers,” J Pineal Res, pp. 253–278, Oct. 2016, doi: 10.1111/JPI.12360.

[18] F. Luchetti et al., “Melatonin Attenuates Ischemic-like Cell Injury by Promoting Autophagosome Maturation via the Sirt1/FoxO1/Rab7 Axis in Hippocampal HT22 Cells and in Organotypic Cultures,” Cells 2022, Vol. 11*, Page* 3701, vol. 11, no. 22, p. 3701, Nov. 2022, doi: 10.3390/CELLS11223701.

[19] L. Dong et al., “Melatonin protects against developmental PBDE-47 neurotoxicity by targeting the AMPK/mitophagy axis,” J Pineal Res, vol. 75, no. 1, p. e12871, Aug. 2023, doi: 10.1111/JPI.12871.

[20] S. Bikri, A. El Mansouri, N. Fath, D. Benloughmari, M. Lamtai, and Y. Aboussaleh, “Melatonin and Hydrogen Sulfide ameliorates cognitive impairments in Alzheimer’s disease rat model exposed to chronic mild stress via attenuation of neuroinflamation and inhibition of oxidative stress: Potential role of BDNF,” Neurosci Behav Physiol, vol. 54, no. 8, pp. 1158–1176, Oct. 2024, doi: 10.1007/S11055-024-01709-4/METRICS.

[21] M. Miranda, J. F. Morici, M. B. Zanoni, and P. Bekinschtein, “Brain-Derived Neurotrophic Factor: A Key Molecule for Memory in the Healthy and the Pathological Brain,” Front Cell Neurosci, vol. 13, Aug. 2019, doi: 10.3389/FNCEL.2019.00363.

[22] E. Castrén and L. M. Monteggia, “Brain-Derived Neurotrophic Factor Signaling in Depression and Antidepressant Action,” Biol Psychiatry, vol. 90, no. 2, pp. 128–136, Jul. 2021, doi: 10.1016/J.BIOPSYCH.2021.05.008.

[23] A. Vohra et al., “Melatonin-mediated corrective changes in gut microbiota of experimentally chronodisrupted C57BL/6J mice,” Chronobiol Int, vol. 41, no. 4, pp. 548–560, 2024, doi: 10.1080/07420528.2024.2329205.

[24] A. Joshi, K. K. Upadhyay, A. Vohra, K. Shirsath, and R. Devkar, “Melatonin induces Nrf2-HO-1 reprogramming and corrections in hepatic core clock oscillations in Non-alcoholic fatty liver disease,” FASEB Journal, vol. 35, no. 9, Sep. 2021, doi: 10.1096/FJ.202002556RRR.

[25] A. H. Vohra, K. K. Upadhyay, A. S. Joshi, H. S. Vyas, J. Thadani, and R. V. Devkar, “Melatonin-primed ADMSCs elicit an efficacious therapeutic response in improving high-fat diet induced non-alcoholic fatty liver disease in C57BL/6J mice,” Egyptian Liver Journal, vol. 11, no. 1, pp. 1–13, 2021.

[26] S. Kulshrestha, R. Karnik, A. Vohra, A. Joshi, and R. Devkar, “Melatonin partially restores hepatic nocturnin oscillations in experimental models of MASLD,” Chronobiol Int, vol. 42, no. 5, pp. 664–677, May 2025, doi: 10.1080/07420528.2025.2496347.

[27] J. Stępniak and M. Karbownik-Lewińska, “Protective Effects of Melatonin against Carcinogen-Induced Oxidative Damage in the Thyroid,” Cancers 2024, Vol. 16, Page 1646, vol. 16, no. 9, p. 1646, Apr. 2024, doi: 10.3390/CANCERS16091646.

[28] G. Ramírez-Rodríguez, N. M. Vega-Rivera, J. Oikawa-Sala, A. Gómez-Sánchez, L. Ortiz-López, and E. Estrada-Camarena, “Melatonin synergizes with citalopram to induce antidepressant-like behavior and to promote hippocampal neurogenesis in adult mice,” J Pineal Res, vol. 56, no. 4, pp. 450–461, May 2014, doi: 10.1111/JPI.12136.

[29] S. Kulshrestha, R. Karnik, A. Vohra, A. Joshi, and R. Devkar, “Exogenous Melatonin Corrects Hepatic Nocturnin Levels in Experimentally Induced MASLD,” Journal of Endocrinology and Reproduction, pp. 103–115, Dec. 2024, doi: 10.18311/JER/2024/44777.

[30] A. K. Gładysz, J. Stępniak, and M. Karbownik-Lewińska, “Exogenous Melatonin Protects against Oxidative Damage to Membrane Lipids Caused by Some Sodium/Iodide Symporter Inhibitors in the Thyroid,” Antioxidants 2023, Vol. 12, Page 1688, vol. 12, no. 9, p. 1688, Aug. 2023, doi: 10.3390/ANTIOX12091688.

[31] S. S. Singh et al., “Melatonin Modulates Hypophyseal-Thyroid Function through Differential Activation of MT1 and MT2 Receptors in Hypothyroid Mice,” Hypothyroidism - New Aspects of an Old Disease, Jan. 2022, doi: 10.5772/INTECHOPEN.100524.

