## Supplementary Files for "Melatonin improves neuro-behavioral perturbations in diet/photoperiod induced chronodisruption"

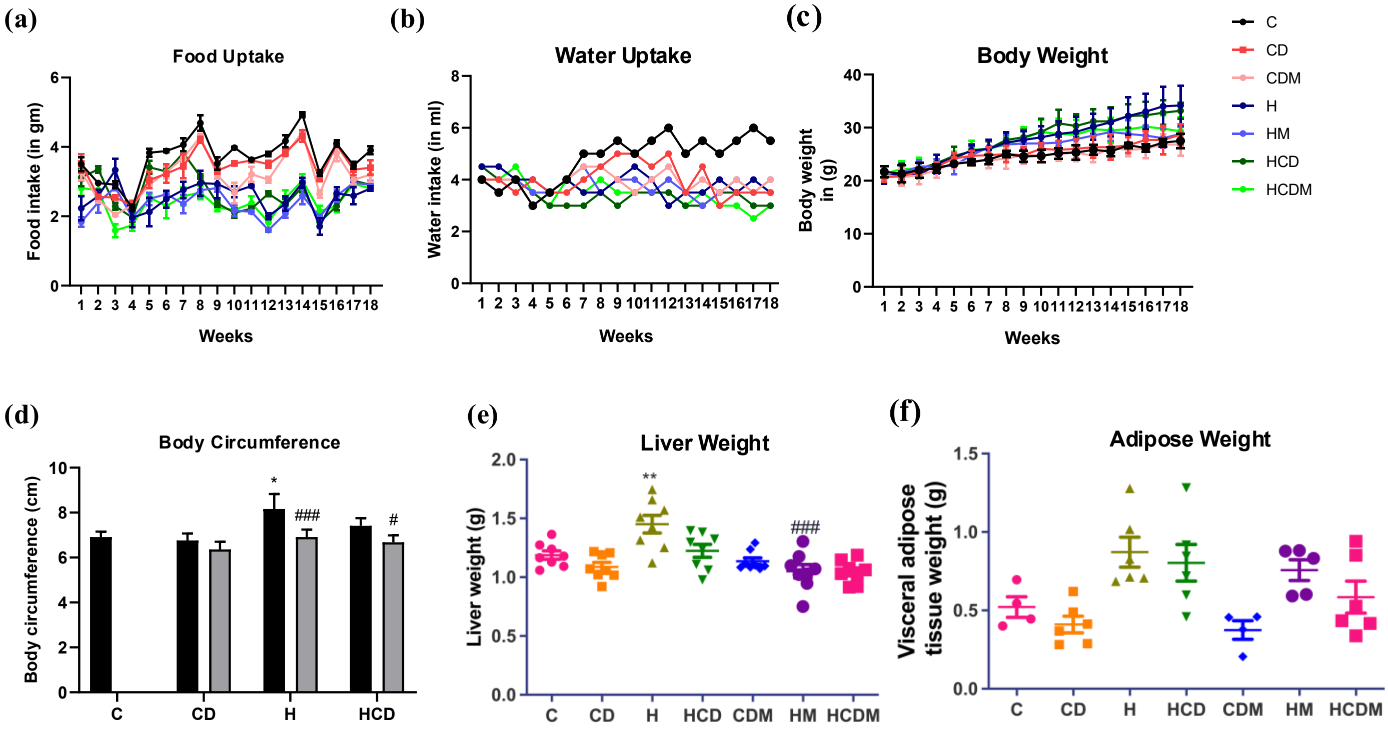


**Supplementary Figure 1:** Changes in (a) Food Intake (gm), (b) Food Intake (ml) and (c) Body Weight (d) Body circumference (cm) (e) Liver Weight (gm) (f) Adipose Weight (gm) of C57BL/6J mice fed with high fat-high fructose diet and/or subjected to chronodisruption and an improvement following melatonin treatment. Results are expressed as mean ± SD *p < 0.05, and ***p < 0.001 is when CD, H and HCD compared to Control (C). #p < 0.05, ##p < 0.01, and ###p < 0.001 is when CDM compared with CD, HM with H and HCDM with HCD respectively.

**Supplementary Table 1:** List of Primers for Real-time PCR

| Gene Name | Forward primer | Reverse primer |
|---|---|---|
| TNF-α | GTGGAACTGGCAGAAGAG | AATGAGAAGAGGCTGAGAC |
| IL-1β | TCTATACCTGTCCTGTGTAATG | GCTTGTGCTCTGCTTGTG |
| IL-4 | GTAGGGCTTCCAAGGTGCTT | GGCATCGAAAAGCCCGAAAG |
| IL-6 | TGGATGCTACCAAACTGGAT | TGGATGCTACCAAACTGGAT |
| IL-10 | AAGGGTTACTTGGGTTGCCA | TTCAGCTTCTCACCCAGGGA |
| IL-12 | ATTACTCCGGACGGTTCACG | ACGCCATTCCACATGTCACT |
| IL-17 | ACCGCAATGAAGACCCTGAT | TCCCTCCGCATTGACACA |
| MCP-1 | TGACCCCAAGAAGGAATGGG | GACCTTAGGGCAGATGCAGTT |
| Nf-κB | GAGGTCTCTGGGGGTACCAT | AAGGCTGCCTGGATCACTTC |
| CREB | GAAGAACAGGGAGGCAGCAA | AGTCCCATTCCTCACACCGT |
| IBA-1 | TGGGTTTGTGTTCGTCAGGC | CATGGTGGGGACAGGAAGTAG |
| BDNF | TCATCCCTCCCCGAGAGTTC | TGGGCTCAATGAAGCATCCG |
| TrkB | GTCAGCCCTCACGTCACTTC | CAACTGCGGTAGCAGGACA |
| SYN-1 | ATCTCTGGTCCCACTCGTCA | ACATCCTGGCTGGGTTTCTG |
| PSD-95 | GAAGACCTCTCAGGCCCCTA | AGGGGTAGGGTTATTGGGCT |
| NT3 | CCGGTGGTAGCCAATAGAACC | GCTGAGGACTTGTCGGTCAC |
| NT-4 | AGCCGGGGAGCAGAGAAG | ACCTCCTCACTCTGGGACTG |
| GAPDH | GTCGGTGTGAACGGATTTGG | TAGATGCCTGCTTCCCATTC |
